# Dissociated cortical phase- and amplitude-coupling patterns in the human brain

**DOI:** 10.1101/485599

**Authors:** Marcus Siems, Markus Siegel

## Abstract

Coupling of neuronal oscillations may reflect and facilitate the communication between neuronal populations. Two primary neuronal coupling modes have been described: phase-coupling and amplitude-coupling. Theoretically, both coupling modes are independent, but so far, their neuronal relationship remains unclear. Here, we combined MEG, source-reconstruction and simulations to systematically compare cortical phase-coupling and amplitude-coupling patterns in the human brain. Importantly, we took into account a critical bias of amplitude-coupling measures due to phase-coupling. We found differences between both coupling modes across a broad frequency range and most of the cortex. Furthermore, by combining empirical measurements and simulations we ruled out that these results were caused by methodological biases, but instead reflected genuine neuronal amplitude coupling. Overall, our results suggest that cortical phase- and amplitude-coupling patterns are non-redundant, which may reflect at least partly distinct neuronal mechanisms. Furthermore, our findings highlight and clarify the compound nature of amplitude coupling measures.

**Highlights:**

- Systematic comparison of cortical phase- and amplitude-coupling patterns
- Demonstration of genuine amplitude coupling independent of phase coupling bias
- Amplitude- and phase coupling patterns differ across many cortical regions and frequencies

## 1. Introduction

The brain is a distributed information processing system. Correlated oscillations of neuronal activity have been proposed to facilitate and orchestrate communication between distant brain regions (Fries, 2015; Siegel et al., 2012; Singer, 1999). In this context, neuronal firing is described as a probabilistic process that is shaped by the phase and amplitude of oscillatory rhythms (Destexhe et al., 1999; Engel et al., 2013, 2001; Fries, 2015; Hillebrand et al., 2012; Hipp et al., 2012; Jahnke et al., 2014; Jensen and Mazaheri, 2010; Siegel et al., 2012). When temporally correlated, co-fluctuations of local oscillations may enhance effective communication between neuronal populations and enable the multiplexing of neuronal information (Akam and Kullmann, 2014; Lopes da Silva, 2013; Singer, 2013). There are two primary coupling modes between neuronal oscillations: phase-coupling and amplitude-coupling (Bruns et al., 2000; Siegel et al., 2012; Engel et al., 2013).

Phase-coupling refers to a consistent phase-alignment between neuronal oscillations, which may reflect a frequency specific signature of neuronal interactions (Siegel et al., 2012). Moreover, phase-coupling may itself modulate effective connectivity by aligning rhythmic excitability fluctuations to rhythmic spike inputs (Fries, 2015). Consistent with this functional role, long-range neuronal phase-coupling reflects various cognitive processes, such as e.g. selective attention (Bosman et al., 2012; Buschman and Miller, 2007; Gregoriou et al., 2009; Siegel et al., 2008), perception (Hipp et al., 2011), memory (Fell and Axmacher, 2011; Palva et al., 2010) and task switching (Buschman et al., 2012). Moreover, task-dependent phase-coupling is expressed in well-structured, large-scale cortical networks (Hipp et al., 2011; Palva et al., 2010).

Amplitude-coupling refers to the temporal co-modulation of the amplitude (or power) of neuronal oscillations. Like phase-coupling, amplitude-coupling may not only result from, and thus reflect, neuronal interactions, but may also regulate these interactions by temporally aligning distant processes associated with fluctuating oscillations (Siegel et al., 2012; von Nicolai et al., 2014). Also amplitude-coupling is expressed in well-structured cortical networks that match known anatomical and functional connectivity (Hipp et al., 2012; Siems et al., 2016), resemble fMRI correlation patterns (Brookes et al., 2011; Deco and Corbetta, 2011; Destexhe et al., 1999; Hipp and Siegel, 2015; Mantini et al., 2007; Nir et al., 2008; O’Neill et al., 2015), and are more stable than phase-coupling networks (Colclough et al., 2016; Wang et al., 2014). Amplitude-coupling is largely driven by amplitude dynamics below 0.1 Hz (Hipp et al., 2012), which may reflect the slow establishment and decay of communicating networks (Destexhe et al., 1999; Leopold et al., 2003; Mantini et al., 2007; Larson-Prior et al., 2011; Hipp et al., 2012; Engel et al., 2013).

Both coupling modes may provide versatile biomarkers for various neuropsychiatric diseases (Fornito et al., 2015; Stam, 2014) including autism (Kitzbichler et al., 2015), schizophrenia (Cetin et al., 2016; Maran et al., 2016), epilepsy (Burns et al., 2014; van Dellen et al., 2014; Zerouali et al., 2016), dementia (Koelewijn et al., 2017; Maestú et al., 2015), Parkinson’s disease (Oswal et al., 2016), multiple sclerosis (Cover et al., 2006; Schoonheim et al., 2013; Tewarie et al., 2014) and blindness (Hawellek et al., 2013). Despite the strong interest and rapidly growing evidence on both, neuronal phase- and amplitude coupling measures, their relationship remains unclear. One the one hand, both coupling-modes could be independent. There could be phase-coupling without amplitude-coupling and vice versa (Siegel et al., 2012). In this case, phase- and amplitude-coupling could be caused by distinct neuronal mechanisms and their cortical coupling-patterns may be dissociated. On the other hand, both coupling modes may be tightly linked, e.g. if both modes reflect the same underlying neuronal interactions, or if one coupling mode causes the other (von Nicolai et al., 2014; Womelsdorf et al., 2007). In this case, the cortical patterns of both coupling modes may be highly similar or even identical. Intermediate scenarios are also possible. The central aim of this study was to non-invasively investigate this relationship between phase- and amplitude coupling in the human brain with MEG.

Addressing this questions is complicated by a methodological peculiarity of the estimation of amplitude coupling that has recently been pointed out (Palva et al., 2018). If erroneous coupling due to field-spread is suppressed by orthogonalization (Brookes et al., 2012; Hipp et al., 2012), measures of amplitude coupling are also partially sensitive to phase coupling (Palva et al., 2018). In other words, the measured amplitude-coupling reflects a mixture of the genuine amplitude-coupling of interest and spurious amplitude-coupling due to phase-coupling.

Thus, we approached our central question in two steps. First, we tested if there is a genuine component to the cortical amplitude-coupling measured with MEG, beyond the spurious amplitude-coupling induced by phase-coupling. Second, we addressed our main question how phase- and genuine amplitude-coupling relate. To this end, we systematically compared the cortical correlation structure of both coupling modes across the human brain.

## 2. Results

We quantified brain-wide neuronal phase- and amplitude-coupling from resting-state MEG measurements in 95 healthy participants. We applied source-reconstruction (Van Veen et al., 1997) to systematically characterize neuronal coupling at the cortical source level. Field spread (or signal leakage) can induce spurious coupling of sensor- and source-level MEG/EEG signals. Thus, we employed two coupling measures discounting signal leakage. For phase-coupling, we applied the weighted phase lag index (wPLI; Nolte et al., 2004; Vinck et al., 2011), which shows the best reliability of volume-conduction free phase-coupling measures (Colclough et al., 2016) and showed the same coupling patterns as the imaginary coherence or the phase lag index (see Fig. S2 for a comparison of the wPLI with these phase-coupling measures). For amplitude coupling, we employed pair-wise signal orthogonalization before estimating amplitude envelope-correlations (Fig. 1A) (Brookes et al., 2012; Hipp et al., 2012).

**Figure 1.**
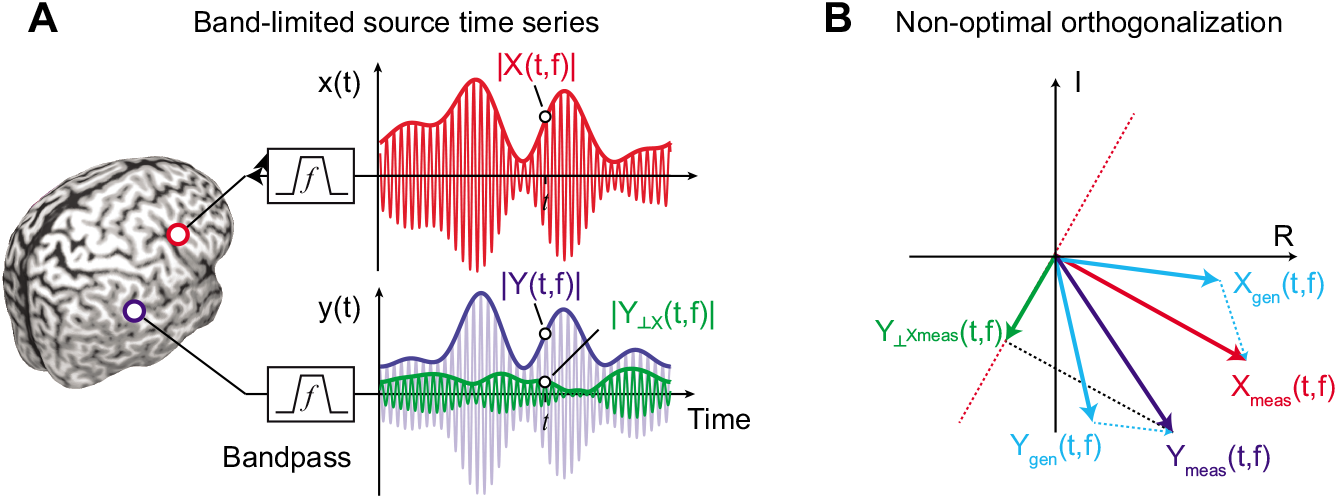
Principle of signal leakage reduction for amplitude relations. (A) Illustration of band-limited time series from two sources X (red, upper panel) and Y (blue, middle panel) with their envelopes (thick lines). The green thick line resembles the envelope of signal Y orthogonalized on signal X. (B) illustrates how the orthogonalization can induce specious amplitude coupling in the presence of phase coupling and signal leakage. We orthogonalize the measures signal Y_meas_ onto the measured signal X_meas_. In the presence of signals leakage, both measured signals reflect a mix of the genuine signals X_gen_ and Y_gen_. For non-zero phase coupling between X_gen_ and Y_gen_, X_meas_ is rotated away from X_gen_. This causes sub-optimal signal orthogonalization.

It has recently been shown that signal orthogonalization does not perfectly discount volume conduction in the presence of genuine phase coupling with non-zero phase delays (Palva et al., 2018). Intuitively, this is because, in the presence of signal leakage, such phase coupling systematically rotates the estimate of the signal to which one aims to orthogonalize, which results in sub-optimal orthogonalization and spurious amplitude-correlations (Fig. 1B). To test if the empirically measured amplitude-coupling patterns reflect this spurious amplitude coupling due to phase coupling, we directly estimated the spurious amplitude coupling with numerical simulations based on empirical parameters (see 2.6). In brief, for each subject, we simulated pairs of cortical signals with no amplitude coupling, with their measured phase coupling (wPLI) at a 90 ° phase shift, and their estimated signal leakage (resolution matrix). With this approach we computed the expected patterns of spurious amplitude coupling under the assumption of no genuine amplitude coupling.

### 2.1 Seed-Based Connectivity Analysis

As a first step, we performed a seed-based analysis (Fig. 2). We computed cortex-wide phase- (Fig. 2C) and amplitude-coupling (Fig. 2A) patterns of neural activity at 16 Hz for several early sensory and higher order cortical regions. As early sensory regions we chose primary auditory (A1) and somatosensory cortex (S1), which show strong inter-hemispheric connectivity and robust amplitude-coupling patterns at 16 Hz (Hipp et al., 2012; Mehrkanoon et al., 2014; Siems et al., 2016). For each seed, subject and both coupling modes, we z-scored the raw coupling measures and tested for z-scores larger than zero across subjects (one-sided t-test, FDR-corrected). This revealed which connections showed significant above-average coupling, discounting global offsets of coupling measures (Hipp et al., 2012).

**Figure 2.**
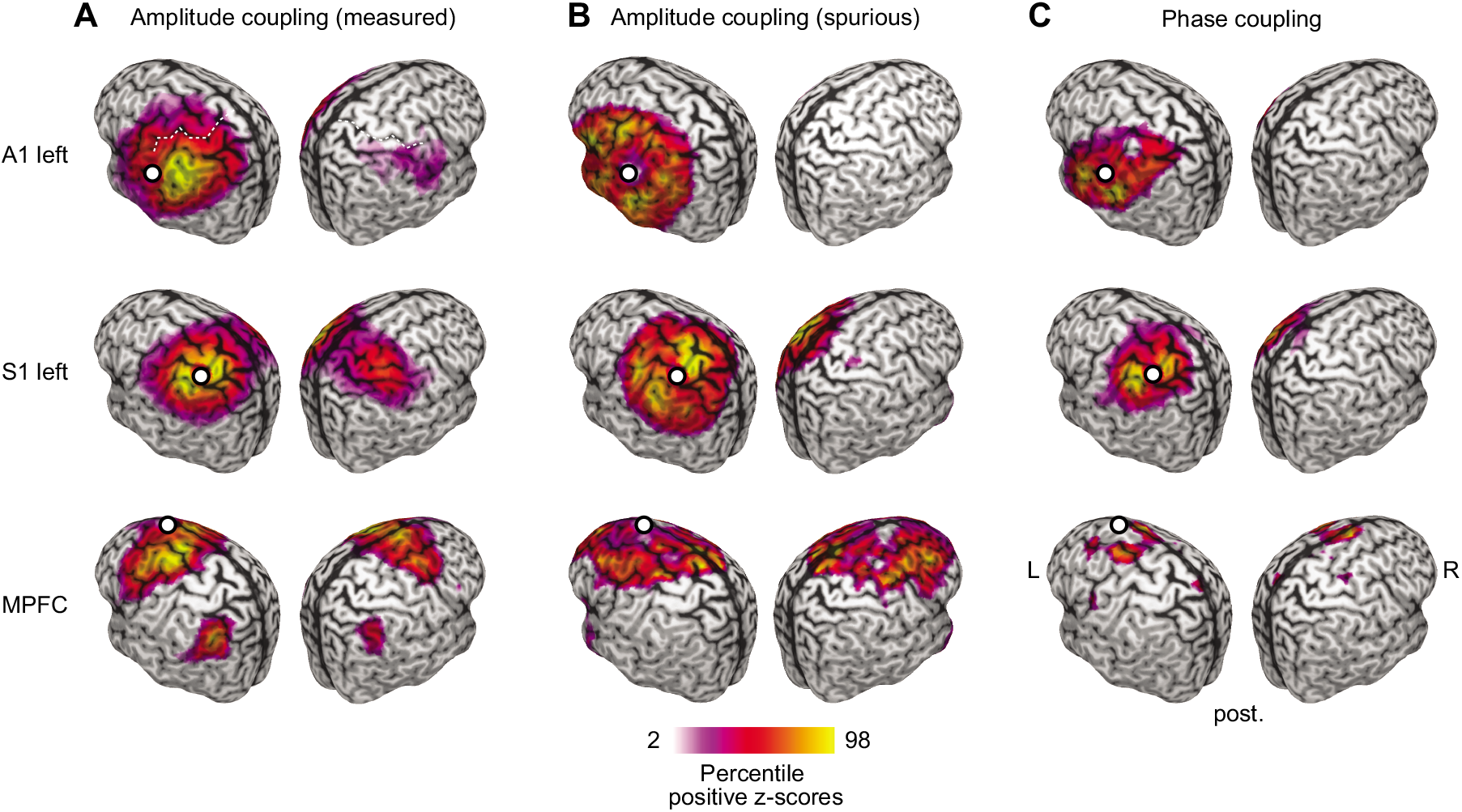
Seed based analysis for early sensory and higher order cortices at 16Hz. Seed-based correlation structure (z-scores) of the left auditory (left A1, top panel), left somatosensory (left S1, middle panel), and the medial prefrontal cortex (MPFC, bottom panel) for measured amplitude-coupling (A), spurious amplitude-coupling due to phase-coupling (B) and phase coupling (C). Coupling z-scores are tested against zero and statistically masked (p<0.05, FDR corrected). Color scale ranges from the 2^nd^ to the 98^th^ percentile of significant values, scaled within each panel. White dots indicate seed regions. The white dashed line in the top left panel highlights the central sulcus (see 4.3 for exact seed coordinates).

For both sensory seeds (A1 and S1), amplitude coupling was strongest to regions surrounding the seed region and to the homologous area in the other hemisphere (Fig. 2 left & right). Phase coupling did not show this pattern, but only above-average connectivity surrounding the seed.

Our findings for a higher order seed region confirmed these results. We investigated phase and amplitude coupling for the medial prefrontal cortex (MPFC, Fig. 2 bottom row), which shows a complex connectivity structure for amplitude coupling at 16 Hz (Hipp et al., 2012; Siems et al., 2016). We found that amplitude coupling of MPFC peaked bilaterally in the dorsal prefrontal and lateral parietal cortices. In contrast, phase coupling only peaked surrounding the seed region.

The spurious amplitude-coupling (Fig. 2B) did not resemble the complex spatial structure of the measured amplitude coupling patterns. In contrast, it showed a stereotypical and robust pattern of above-average connectivity only to sources near the seed, decreasing with distance independent of the seed location.

### 2.2 Genuine amplitude coupling

To quantitatively address our first question, i.e. if the measured amplitude coupling (AC_meas_) reflects genuine amplitude coupling, we systematically assessed the similarity of the cortical patterns of spurious (P_ACspur_) and measured amplitude coupling (P_ACmeas_) across frequencies (Fig. 3). A high correlation would indicate that the measured amplitude coupling could largely be explained by spurious amplitude-coupling. Conversely, the fraction of non-explained amplitude coupling will be attributed to genuine amplitude coupling.

**Figure 3.**
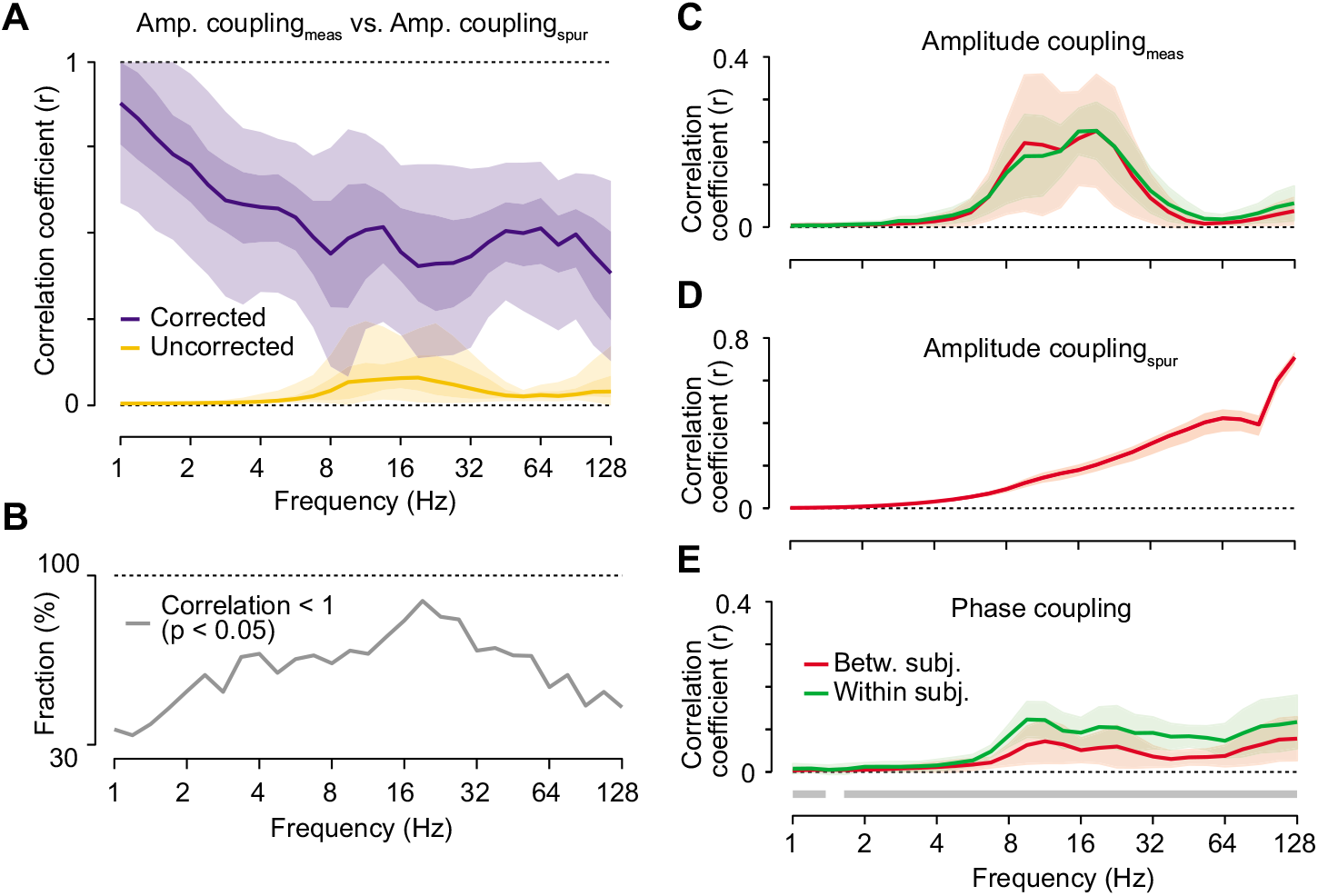
Correlation between measured and spurious amplitude-coupling patterns. (A) Frequency resolved correlation between measured and spurious amplitude-coupling patterns. Lines indicate median attenuation corrected (blue) and uncorrected (yellow) correlation. Shaded areas indicate the 5-95% and 25-75% inter-percentile range over space. (B) Fraction of cortical locations that show significant (p<0.05, FDR-corrected) genuine amplitude coupling, i.e. a correlation between measured and spurious amplitude coupling, corrected for pattern reliabilities, is significantly smaller than 1 (see 4.9). (C) Reliability, i.e. correlation, of measured amplitude-coupling patterns within (green) and between (red) subjects. Shaded areas indicate the 25-75% interquartile range. (D) Reliability of spurious amplitude-coupling patterns between subjects. (E) Reliability of phase-coupling patterns within (green) and between (red) subjects. The gray bars below the lines indicates significantly larger reliability within as compare to between subjects (p<0.05, FDR-corrected).

For each frequency and both measures, we computed the coupling between all cortical regions, i.e. we computed the full connectivity matrices of the cortex-wide measured and spurious amplitude coupling. We then correlated the patterns of spurious and measured amplitude coupling for each cortical seed region (3 examples from the 457 sources in one frequency are shown in Fig. 2). In other words, we correlated each column of the connectivity matrices between measures. Averaged across all seed regions, this revealed a positive correlation that markedly peaked from 8-32 Hz and above 90 Hz with median correlation coefficients below 0.1 (Fig. 3A, yellow line).

At first sight, the low correlation between measured and spurious amplitude-coupling patterns suggests that there is indeed genuine amplitude coupling. However, it is important to realize that the correlation between two metrics does not only reflect their true underlying correlation, but also the metrics’ reliability (Bergholm et al., 2010; Hipp et al., 2012; Siems et al., 2016; Spearman, 1904). A low reliability of two measures, e.g. due to strong noise, leads to a low measured correlation even if the true underlying correlation between the two measures is high (Supplementary Fig. 1, dashed lines). Thus, the observed low and frequency specific correlation between spurious and measured amplitude-coupling patterns may merely reflect the low and frequency specific reliability of either measure, and thus, does not allow for directly inferring genuine amplitude coupling.

We applied attenuation correction of correlations (Hipp and Siegel, 2015; Siems et al., 2016; Spearman, 1904) to account for the effect of signal reliability. Attenuation corrected correlations quantify how strong a correlation would be for perfectly reliable signals (Fig. S1). We employed the between-subject correlation of the measured and spurious amplitude coupling-patterns as a proxy for each measure’s reliability (Hipp and Siegel, 2015; Siems et al., 2016). For the measured amplitude coupling, between-subject reliability peaked around 16 Hz (Fig. 3C) compatible with previous findings (Hipp and Siegel, 2015; Siems et al., 2016). For the spurious amplitude-coupling, reliability increased monotonically with frequency (Fig. 3D), which likely reflects the decreasing spatial resolution of beamforming filters with increasing frequency.

We corrected the correlation between measured and spurious amplitude-coupling patterns for these reliabilities by pooled division (see 4.8). This correction had a marked effect (Fig. 3A, blue line). As predicted, the overall correlation between measured and spurious amplitude-coupling patterns increased. The median attenuation corrected correlation was around 0.5 for frequencies above 5Hz and further increased for lower frequencies. This suggests that, for frequencies above 5 Hz, on average more than 70 % of the variance in the measured amplitude-coupling patterns was due to genuine amplitude coupling. This result was largely independent of the phase shift between signals employed for estimating the spurious amplitude-coupling. Also for phase shifts smaller than 90 °, across all investigated frequencies, the attenuation corrected correlation between measured and spurious amplitude-coupling patterns was substantially smaller than 1 (Fig. S3).

The positive correlation between spurious and measured amplitude-coupling patterns may not only reflect non-optimal orthogonalization, but also a true similarity between these patterns. Nearby sites may show genuinely higher amplitude correlations similar to the pattern of the spurious amplitude-coupling that reflects nearby field-spread and local phase-coupling (compare Fig. 2C). In contrast, more complex patterns in amplitude coupling display a non-monotonous distance relation (for example Fig. 2 MPFC). To investigate the dependency on cortical distance, we split each seed correlation-pattern in four distance quartiles and repeated our analysis separately, for each quartile (Fig. 4). As hypothesized, we found that the similarity between spurious and measured amplitude-coupling patterns was highest for the closest connections. For longer distances, the correlation decreased indicating a stronger dissimilarity between spurious and measured amplitude-coupling patterns.

**Figure 4.**
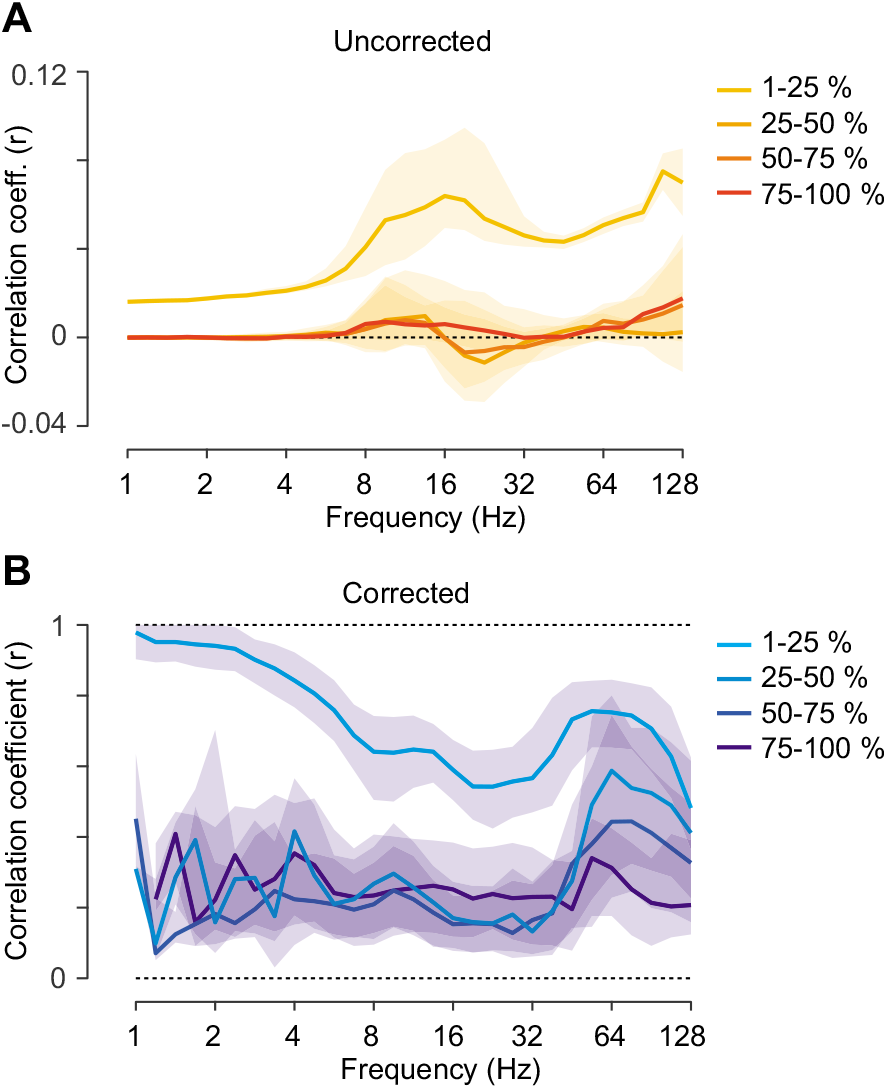
Correlation between measured and spurious amplitude-coupling patterns for separate coupling distances. (A) Frequency resolved un-corrected correlation between measured and spurious amplitude-coupling patterns for four coupling-distance quartiles. For the correlation between both measures, the patterns (columns of the correlation matrix) were split into four quartiles depending on the distance of each source from the seed. Lines indicate median correlation. Shaded areas indicate the 25-75% interquartile range over space. (B) Frequency resolved attenuation-corrected correlation between measured and spurious amplitude-coupling patterns for four coupling-distance quartiles.

We next sought to statistically assess if there was any genuine amplitude coupling, i.e. if the attenuation corrected correlations between measured and spurious amplitude-coupling patterns were indeed significantly smaller than 1. Attenuation corrected correlation is an unbiased estimate (Figure S1). Thus, we applied a leave-one-out jackknifing procedure and false-discovery rate correction (Benjamini and Hochberg, 1995). Across the entire spectrum, we found that on average more than 50% of the seed patterns showed significant (p < 0.05 corrected) genuine amplitude coupling (Fig. 3B). The amount of significant differences peaked around 22Hz with more than 90% of seed patterns with a correlation significantly smaller than 1. It should be noted that, in contrast to the attenuation corrected correlation itself, this statistic is confounded by the measurement reliability. Areas with high reliability show less variability across subjects, and hence, increased statistical power. Thus, the attenuation corrected correlation itself (Fig. 3A), rather than its statistic, should be used to quantitatively assess the dependency between measured and spurious amplitude-coupling patterns.

The cortical distribution of these attenuation corrected correlations indicates, which areas show the lowest dependency between measured and spurious amplitude coupling patterns, and thus, potentially the strongest genuine amplitude coupling (Fig. 5A, see Figure S5 for statistical masking). This cortical distribution showed a complex pattern across frequencies. A cluster analysis revealed four major patterns (Fig. 5B and C, lowest BIC for 4 clusters):

**Figure 5.**
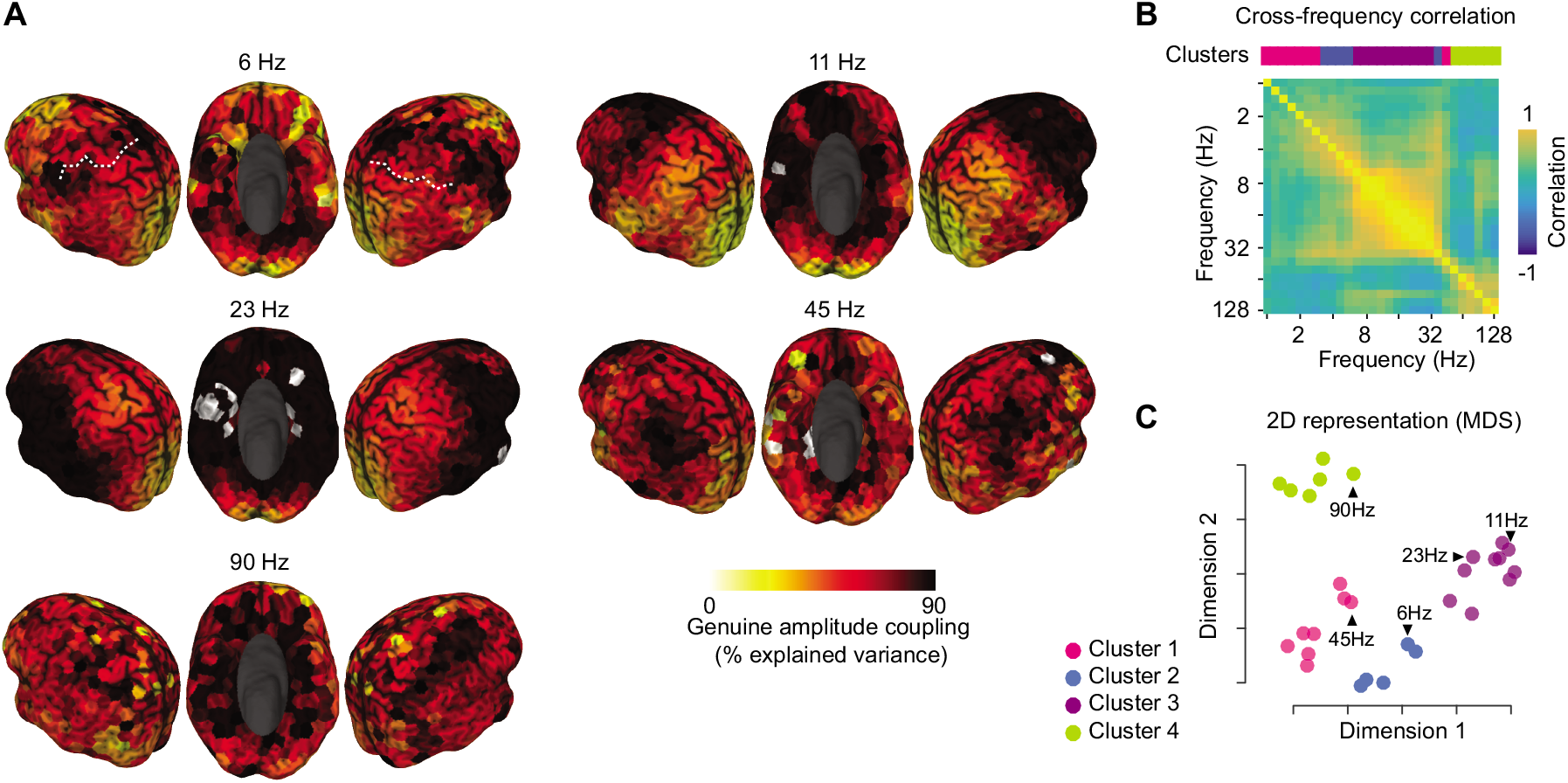
Spatial distribution of genuine amplitude coupling. (A) Spectrally and spatially resolved fraction of variance in the measured amplitude coupling that cannot be explained by spurious amplitude-coupling due to phase-coupling and is thus attributed to genuine amplitude coupling (attenuation corrected). The color scale indicates the amount of genuine variance, i.e. 1-r^2^ of the attenuation corrected correlation between measured and spurious amplitude-coupling. The white dashed line in the top left panel indicates the central sulcus. See Figure S5 for statistical masking. (B) Cross-frequency correlation of attenuation corrected correlation patterns. The colored bar on top of the matrix indicates the frequency specific clustering of patterns (Gaussian mixture model with 4 clusters for minimal BIC, see 4.11). (C) 2D multidimensional scaling representation of cortical patterns in (A) based on the Euclidean distance between patterns (see 4.11). Points are colored according to the clustering of patterns.

From about 4 to 32 Hz (cluster 2 & 3) genuine amplitude coupling peaked in the lateral and posterior prefrontal cortex and around the temporal pole (Fig. 5A & S4A). Cluster 3 (7-27 Hz) showed peak genuine amplitude coupling in medial prefrontal and orbitofrontal cortex (Fig. 5A & S4A). For low frequencies (1-3 Hz) and around 45 Hz, patterns appeared scattered with strongest genuine amplitude coupling ventrolateral prefrontal areas. For high frequencies (> 64 Hz, cluster 4), genuine amplitude coupling primarily peaked in basal and lateral parietal areas. For all investigated frequencies, early visual areas showed the strongest similarity between measured and spurious amplitude-coupling patterns.

In sum, these results indicate that the measured amplitude coupling is indeed different from the spurious coupling introduced by phase coupling and volume conduction. Furthermore, for most of the brain and frequencies, genuine amplitude coupling likely accounts for 70 % and more of the variance in cortical amplitude coupling patterns.

### 2.3 Comparing genuine amplitude-coupling and phase-coupling networks

Establishing a genuine component in amplitude-coupling patterns, allowed us to address our second question, i.e. if there are differences between amplitude- and phase-coupling patterns. Although attenuation correction estimates the amount of variance due to genuine amplitude coupling, we cannot directly extract genuine amplitude-coupling patterns. This precludes a direct comparison between genuine amplitude-coupling and phase-coupling patterns. However, we can indirectly infer differences between these patterns using a geometric heuristic. For this, we conceptualize the connectivity patterns as vectors within a high-dimensional space (Fig. 6). Here, their co-linearity describes their similarity, i.e. correlation. We assume that the spurious amplitude-coupling patterns (P_ACspur_) are a positively weighted sum of the phase coupling (P_PC_) and field spread (P_M_) patterns (Fig. 6A). Thus, P_ACspur_ is situated in the hyper-area between P_PC_ and P_M_. Under the Null-hypothesis that we want to test, the genuine amplitude-coupling pattern (P_ACgen_) is identical to the phase-coupling pattern (Fig. 6B). As the measured amplitude coupling pattern (P_ACmeas_) is assumed to be a positively weighted summation of the spurious and genuine amplitude coupling patterns, under the Null hypothesis, P_ACmeas_ is situated in the hyper-area between the spurious amplitude-coupling patterns and the phase-coupling patterns (Fig. 6B). Hence, if the measured amplitude coupling is outside this Null-hypothesis area, we accept the alternative hypothesis that genuine amplitude-coupling patterns and phase-coupling patterns are not identical (Fig. 6C). P_ASmeas_ is outside the Null-hypothesis area either if the PACspur-PPC correlation is stronger than the PACmeas-PPC correlation (condition 1) or if the PACspur-PPC correlation is stronger than the PACmeas-PACspur (condition 2) (Fig. 6C). To test these conditions, we computed attenuation corrected correlations between PACspur-PPC, PACmeas-PPC and PACmeas-PACspur, and applied leave-one-out Jackknifing to test for significant differences (FDR-corrected).

**Figure 6.**
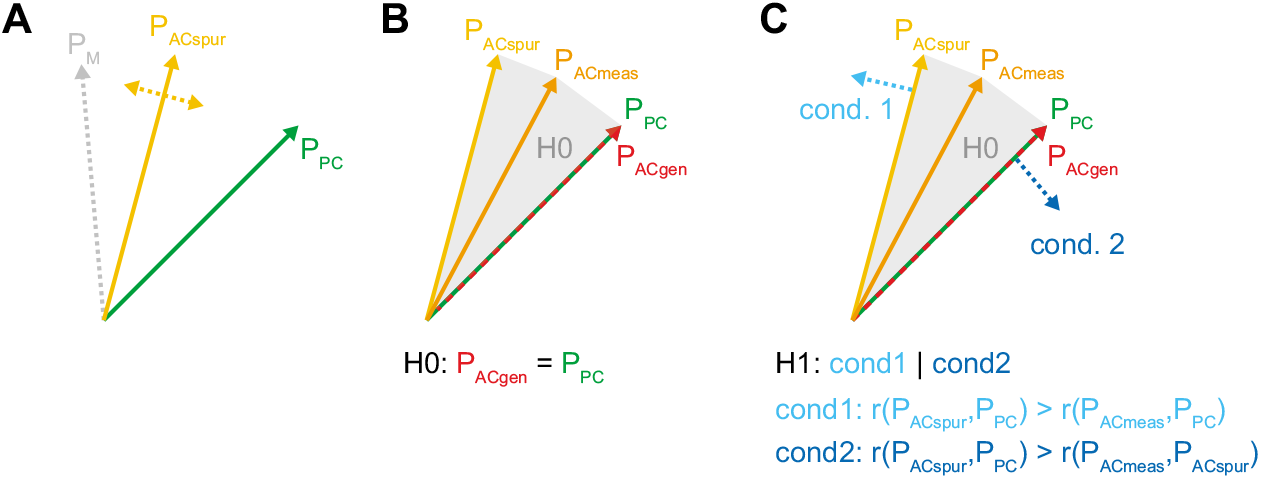
Schematic comparison of phase-coupling and genuine amplitude-coupling patterns. Different phase-coupling and genuine amplitude-coupling patterns can be inferred if either one of two critical conditions are met. This is illustrated here by conceptualizing the cortical patterns, which for the present data are 457-dimensional vectors, as two-dimensional vectors with unit length. The co-linearity between vectors corresponds to the correlation of these patterns. (A) The mixing pattern (PM) (field spread) and the phase coupling pattern (PPC) are empirically measured. The pattern of the spurious amplitude-coupling (PACspur) corresponds to a weighted average of PPC and PM. (B) The measured pattern of amplitude coupling (PACmeas) is a weighted average of the pattern of genuine amplitude coupling (PACgen) and the spurious amplitude-coupling (PACspur). Under the Null-hypothesis that genuine amplitude coupling and phase coupling patterns are identical, P_ACmeas_ is situated in the hyper-area between P_ACspur_ and P_PC_ (H0-area). (C) If P_ACmeas_ is outside the H0-area, we accept the alternative hypothesis that P_ACgen_ and P_PC_ are different. P_ACgen_ is outside the H0-area if condition 1 (light blue) or condition 2 (dark blue) is met.

We found that, for large parts of the brain and for all frequencies, there were significant differences between cortical phase- and amplitude-coupling patterns (Fig. 7). For all frequencies, amplitude and phase coupling differed for at least 50% and up to 80% of the cortex with significant genuine amplitude coupling patterns (Fig. 7B).

**Figure 7.**
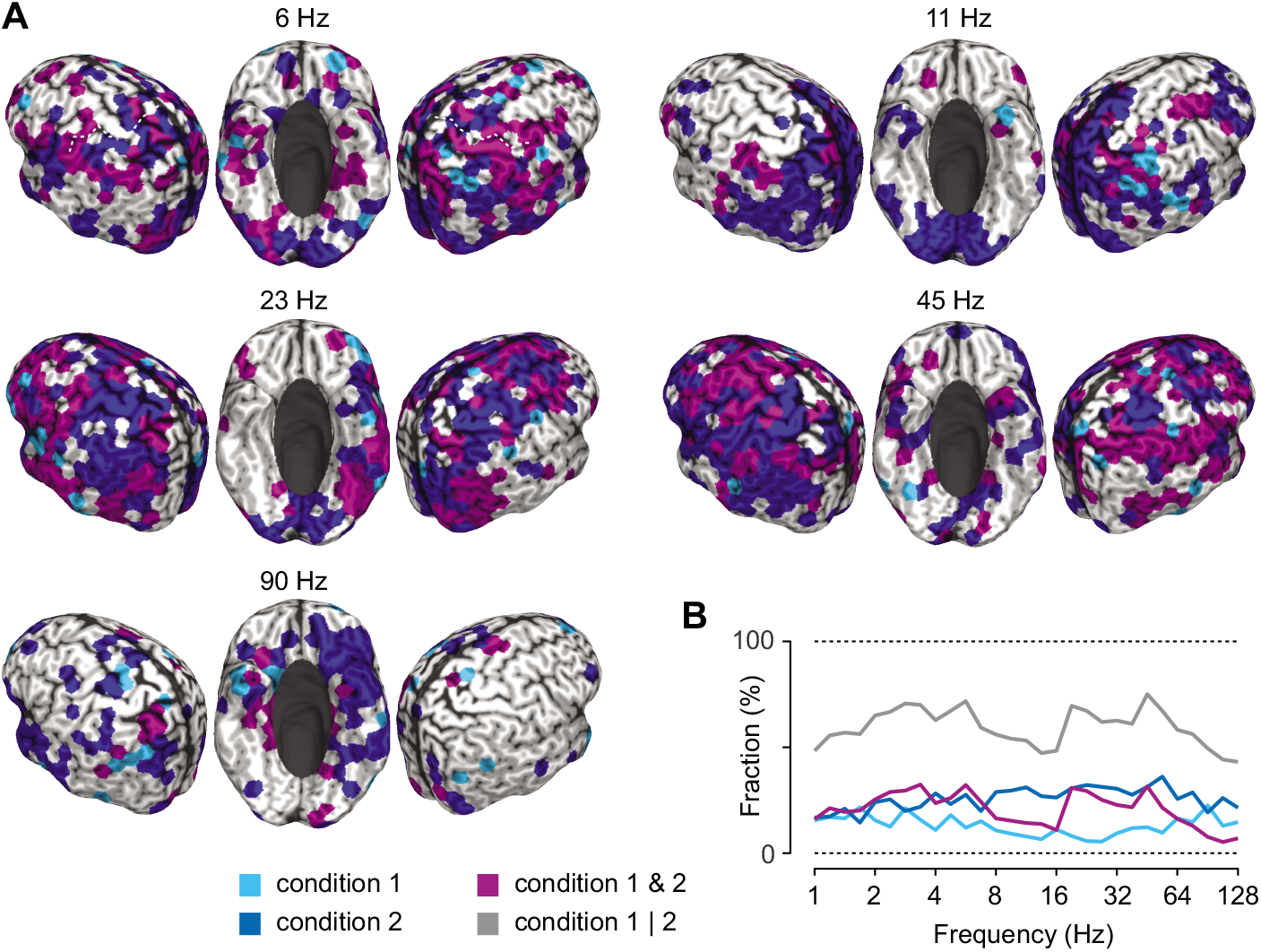
Differences between genuine amplitude-coupling patterns and phase-coupling patterns. (A) Cortical distribution of regions that show significant differences between genuine amplitude-coupling and phase-coupling patterns by fulfilling either condition 1 (light blue), condition 2 (dark blue) or both (purple). (B) Fraction of cortical regions that show significant differences between genuine amplitude-coupling and phase-coupling patterns. Same color-scheme as in (A). In addition, the gray line indicates the fraction of regions that fulfill either condition 1 or 2 (see 4.10 for details).

These results establish a clear distinction between amplitude- and phase coupling on the network level. The dissociation between conditions 1 and 2 provided further insights into the nature of this distinction. For almost all carrier frequencies above 2 Hz, condition 2 was met more often than condition 1 alone (Fig. 7A and B). For condition 2, the genuine amplitude-coupling pattern (P_ACgen_) is further away from the field-spread pattern (P_M_) than the phase coupling pattern (P_PC_). Thus, in areas were only condition 2 is met, more complex amplitude coupling patterns may predominate. This included bilateral extrastriate visual areas around 11 Hz, bilateral parietal and pericentral areas at 22 Hz, and the left pericentral and posterior temporal areas at 45 Hz. Overall, our results indicate a wide-spread dissociation between genuine amplitude- and phase coupling patterns.

## 3. Discussion

Our results provide, to our knowledge, the first systematic comparison of cortical phase- and amplitude-coupling patterns in the human brain. We found differences between both coupling modes that are widely distributed across frequencies and the entire cortex. By combining empirical measurements and simulations we show that these differences are not caused by known methodological biases, but instead reflect a genuine neuronal dissociation. The observed differences suggest that cortical phase- and amplitude-coupling patterns do not solely display redundant network information. This suggests that distinct neural mechanisms at least partly underlie these two coupling modes in the human brain. Furthermore, our results highlight and clarify the compound nature of amplitude coupling measures applied to orthogonalized signals.

### 3.1 Discounting confounding factors

Our analyses discount two critical factors that confound the estimation of neuronal coupling patterns and their comparison. First, we employed amplitude correlations of orthogonalized signals (Brookes et al., 2012; Hipp et al., 2012) and the weighted phase-lag index (Vinck et al., 2011). Using these coupling measures ensured that the measured coupling did not reflect spurious coupling due to field-spread.

Second, for the comparison between coupling modes, we employed attenuation correction of correlations (Spearman, 1904). This approach allows to correct for the attenuation of measured correlation caused by sub-optimal measurement reliability. Attenuation correction of correlations is a powerful analytical approach that has been successfully employed before to compare MEG with fMRI (Hipp and Siegel, 2015) and MEG with EEG (Siems et al., 2016). Importantly, reliability in the present study refers to the stability of coupling patterns across subjects, which effectively takes into account all sources of variance across subjects, including measurement and finite-sampling noise, noise caused by neural activity not of interest, and inter-subject variability. The employed approach corrects for all these sources of variance, which attenuate measured correlations and may thus induce spurious spectral and spatial specificity.

Indeed, our results indicate that the raw correlation between measured and spurious amplitude-coupling patterns is strongly affected by measurement reliability. Attenuation correction suggests that the peaked raw correlation around 16 Hz merely reflects the strength of intrinsic cortical rhythms around this frequency, rather than a frequency specific relation of the measured and spurious amplitude-coupling patterns. The same arguments hold for the comparison between amplitude- and phase-coupling (Zhigalov et al., 2017).

### 3.2 Phase-coupling sensitivity of orthogonalized amplitude correlation

Our results provide a critical reassessment of well-established amplitude-coupling measures of orthogonalized signals (Brookes et al., 2012; Hipp et al., 2012). It has recently been pointed out that, in the presence of field-spread, these measures are sensitive to phase coupling with non-zero phase lag (Palva et al., 2018). Here, we combined the simulation approach put forward by Palva and colleagues (2018) with empirical measurements to systematically evaluate the sensitivity of these measures to phase coupling across the human cortex.

The extent to which phase coupling can induce spurious amplitude-coupling measures critically depends on the amount of field-spread. In agreement with this notion, we found that in particular short distance connections show spurious amplitude coupling (Fig. 2 & Fig. 4). On a brain wide scale, this appears as a stereotypical pattern of connectivity decreasing with distance from the seed regions, independent of seed location or frequency.

### 3.3 Determining genuine amplitude coupling

The bias of orthogonalized amplitude coupling measures left open the possibility that the described amplitude coupling patterns (Brookes et al., 2012; Hipp et al., 2012) merely reflect phase coupling in combination with field spread. Our results provide several lines of evidence against this hypothesis.

First, the spurious amplitude-correlation showed a stereotypical pattern of connectivity decreasing with distance, whereas the measured amplitude-coupling patterns showed complex and multimodal distributions (Fig. 2). The distance-resolved comparison further supports the notion of more complex measured amplitude coupling patterns than expected for a mere bias (Fig. 4). Second, the between subject reliability of coupling patterns clearly dissociated measured and spurious amplitude coupling (Fig. 3C and D). Third, for a large portion of the cortex and frequencies, the correlation of spurious and measured amplitude-coupling patterns was significantly smaller than 1 (Fig. 3B). On average, only 30 % of the variance in amplitude coupling patterns could be explained by spurious amplitude-coupling (only 10% for long connections, Fig. 4B). Finally, the correlation of spurious and measured amplitude-coupling patterns showed a frequency specific cortical distribution (Fig. 5), which is not expected for a stereotypic measurement bias.

In summary, our results show that the amplitude correlation of orthogonalized signals is indeed a compound measure of connectivity, that, in particular for short distances, also reflects phase-coupling. Nevertheless, our results suggest that, beyond spurious amplitude-coupling induced by phase-coupling, the observed amplitude coupling patterns are to a large extent driven by genuine amplitude coupling.

### 3.4 Relation between phase- and amplitude coupling

Which factors may cause the observed differences between phase- and amplitude coupling patterns (Fig. 7)?

First, different non-linearities between coupling modes may induce differences. The same underlying neuronal interaction or common input may have different effects on both coupling modes, thus reducing their correlation. However, in contrast to our present results this effect should be spectrally and spatially unspecific. Thus, such non-linearities cannot entirely explain our findings.

Second, phase- and amplitude-coupling may be differentially affected by non-neuronal artifacts. A particularly strong artifact for frequencies above about 30 Hz is muscle activity. Even if M/EEG data is preprocessed to minimize muscle artifacts, as done here (Larson-Prior et al., 2013), residual muscle activity is often detectable in frontal and temporal regions (Hipp and Siegel, 2013; Siems et al., 2016). Notably, we found strong differences between phase- and amplitude coupling at very high frequencies in these regions (Fig. 7A). This suggests that the two coupling modes may be differentially susceptible to muscle activity. Indeed, specifically the reliability of phase coupling increased at high frequencies (Fig. 3E), which may further indicate stronger susceptibility to muscle artifacts for this coupling mode.

Third, imperfections of the employed simulations and thus of the estimated spurious amplitude-coupling may lead to an underestimation of the similarity of phase and genuine amplitude-coupling patterns (see also 3.6 below).

Finally, distinct neuronal mechanisms may underlie both coupling modes. On the one hand, for example, neuromodulation may co-modulate the strength of rhythms in different brain regions. Or, as recently proposed, slow fluctuations of extracellular potassium concentrations and structural connectivity may drive long-range power cofluctuations (Krishnan et al., 2018). These mechanisms may induce amplitude coupling on a slow temporal scale without necessarily causing phase coupling on a fast temporal scale. On the other hand, synaptic interactions triggered by intrinsic activity or sensory inputs may induce phase-coupling between areas without driving identical amplitude comodulations.

Despite the observed significant differences between coupling modes, our results are also compatible with similarities between their cortical patterns (compare Fig. 2). Such similarities may result from one or more common underlying neural mechanism. Synaptic interactions between neuronal populations may induce both, coupling of phases and amplitudes of these neuronal populations. Similarly, common input to neuronal populations will co-modulate and thus couple both, phases and amplitudes (Tewarie et al., 2018). Alternatively, also causal relations between both coupling modes may result in correlations. For example, as discussed next, phase-locking may enhance neuronal interactions, and thereby, enhance amplitude coupling (Fries, 2015; Womelsdorf et al., 2007).

### 3.5 Functional role of coupling modes

Phase-coupling of neuronal population may regulate their interactions by aligning rhythmic excitability fluctuations and rhythmic inputs (Fries, 2015). Similarly, amplitude-coupling may modulate interactions by temporally aligning processing associated with low or high oscillatory amplitudes across brain regions (Siegel et al., 2012; von Nicolai et al., 2014). While the observed differences between coupling modes may reflect such functional roles, the present results hold independent from such potential functions. In fact, even if phase- or amplitude coupling merely reflect neural interactions without a causal mechanistic role, our results show that these coupling modes provide partially dissociated and thus non-redundant information about neuronal interactions. This suggests that both coupling modes provide complimentary information on large-scale neuronal interactions during cognitive processes and on their alteration in neuropsychiatric diseases.

### 3.6 Limitations

Our results are based on specific parameter choices for the simulations used to estimate the spurious amplitude-coupling. This entails in particular a constant phase-shift and Gaussian distribution of cortical signals. Deviations from these assumptions will lead to sub-optimal estimation of both, the spurious and genuine amplitude coupling patterns. Non-constant phase-shifts may induce patterns of the spurious amplitude coupling that are not reflected in the present simulation. The signal distribution may have an effect on two levels. First, non-Gaussian signals will lead to sub-optimal orthogonalization (Brookes et al., 2014, 2012; Hipp et al., 2012). Second, the spurious amplitude coupling of non-Gaussian signals will deviate from the simulated estimates based on Gaussian signals. Thus, optimal estimation of the spurious and genuine amplitude coupling requires reliable empirical assessment of cortical phase-shift, likely using invasive approaches, and a further systematic assessment of the effect of signal distributions.

### 3.7 Future directions

Our results provide a critical first step to unravel the relationship between neuronal phase- and amplitude-coupling. Further invasive studies are needed to investigate this relationship and the underlying mechanisms on the cellular and circuit level, as well as to link the present results to spiking activity of individual neurons. Additionally, the investigation of non-linear and cross-frequency relationships, i.e. between phase- and amplitude-coupling across different frequencies (Brookes et al., 2016; Diekelmann and Born, 2010; Mandke et al., 2018; Schroeder and Lakatos, 2009; Tewarie et al., 2016; von Nicolai et al., 2014; Womelsdorf et al., 2007) as well as the application of directed interaction measures (Hillebrand et al., 2016; Lobier et al., 2014; Vinck et al., 2015) may allow identifying generic links between coupling modes.

## 4. Methods and Materials

### 4.1 Subjects and dataset

We analyzed resting-state MEG measurements from 95 subjects included in the publicly available human connectome project (HCP) S900 release. Participants were healthy adults in the age range between 22-35 (n_22-25_ = 18, n_26-30_ = 40, n_31-35_ = 37). The sample included 45 females. The resting-state measurements included three six-minute blocks with short breaks in between measurements. Data were recorded with a whole-head Magnes 3600 scanner (4D Neuroimaging, San Diego, CA, USA) situated in a magnetically shielded room (for further details see: Larson-Prior et al., 2013). Additionally, subjects were scanned on a Siemens 3T Skyra to acquire structural T1-weighted magnetic resonance images (MRI) with 0.7mm isotropic resolution (Van Essen et al., 2013).

### 4.2 Data preprocessing

We used the preprocessed data as provided by the HCP pipeline (Larson-Prior et al., 2013). This includes removal of noisy and bad channels, bad data segments and physiological artifacts by the iterative application of temporal and spatial independent component analysis (ICA) (Larson-Prior et al., 2013; Mantini et al., 2011).

### 4.3 Physical forward model and source modeling

MEG sensors were aligned to the individual anatomy using FieldTrip (Oostenveld et al., 2010). We segmented the individual T1-weighted images and generated a single shell head model to compute the physical forward model (Nolte, 2003). We computed the forward model for 457 equally spaced (~1.2cm distance) source points spanning the cortex at 0.7 cm depth below the pial surface (Hipp and Siegel, 2015). This source shell was generated in MNI-space and non-linearly transformed to individual headspace. Source coordinates, head model and MEG channels were co-registered on the basis of three head localization coils.

The sensor-level MEG data was projected to source space using linear beamforming (Gross et al., 2001; Van Veen et al., 1997). This spatial filtering approach reconstructs activity of the sources of interest with unit gain while maximally suppressing contributions from other sources.

Coordinates for the seed-based connectivity analyses were adopted from Hipp et al. (2012). For every seed, the source location of the 457 shell positions with minimum Euclidean distance from the seed coordinates was chosen: left auditory cortex (lAC) [−54, −22, 10]; left somatosensory cortex (lSSC) [42, −26, 54]; medial prefrontal cortex (MPFC) [−3, 39, −2] (all MNI coordinates).

### 4.4 Spectral analysis

Time-frequency estimates of the time-domain MEG signal were generated using Morlet’s wavelets (Goupillaud et al., 1984). The bandwidth of the wavelets was set to 0.5 octaves (1 spectral standard deviation) with a temporal step-size of half the temporal standard deviation. We derived spectral estimates for frequencies from 1 to 128 Hz in quarter octave steps.

### 4.5 Coupling measures

We estimated amplitude coupling using amplitude envelope correlations of orthogonalized signals (Hipp et al., 2012). Volume conduction effect were discounted by orthogonalizing the two complex signals at each point in time before correlation (Brookes et al., 2012; Hipp et al., 2012).

Here, the *imag* operator describes the imaginary part of the signal. The complex signals *x* and *y* are a function of time and frequency. *x’* is the complex conjugate of signal *x*. Discounting volume conduction with orthogonalization is only optimal for data with a Gaussian distribution (Brookes et al., 2014). Finally, we computed the Pearson correlation between the logarithm of power envelopes of the signals *x* and *y_orth_*.

As a measure of phase coupling we applied the weighted phase lag index (wPLI; Vinck et al., 2011). The wPLI takes only the imaginary part of the cross-spectrum into account and normalizes it with the average absolute imaginary contribution within the time series.

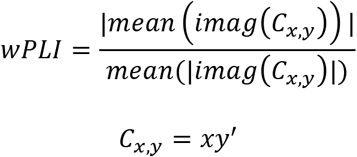

Here, *C_x,y_* is the cross-spectrum between the two complex signals *x* and *y* defined as the product of *x* and the complex conjugate of *y*. The imaginary part of the cross-spectrum is insensitive to volume conduction since it has no contribution from zero phase lagged parts of the signal (Nolte et al., 2004; Vinck et al., 2011). We computed both coupling measures for the full correlation matrices for all subjects and frequency bands.

### 4.6 Data simulation

Palva and colleagues (2018) showed that amplitude correlations based on orthogonalized signals yield spurious correlations, given a consistent non-zero phase delay between signals. We employed the simulation approach put forward by Palva and colleagues (2018) as a generative model to estimate these spurious correlations. We computed a model for every connection, subject and frequency using empirical values for the free parameters. With this approach, we generated complete correlation matrices for every subject and frequency to estimate the spatial patterns of spurious amplitude-coupling. We modeled every two signals *x* and *y*

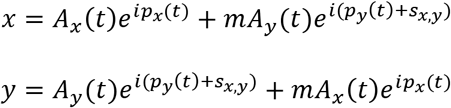

where *A(t)* and *p(t)* are vectors representing the amplitude and the phase of the sources, respectively. In analogy to volume conduction, the source data is linearly mixed by the parameter *m*. This value is determined from the empirical data as the multiplication of the filter matrix *F_x,f_* with the leadfield *L_y_* (the resolution matrix) projected onto the first principal dipole direction *P1* at *x*.

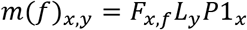

For every connection, we computed the model in both directions. We averaged the two directions, *x* on *y* and *y* on *x*, in the final correlation matrices. *s_x,y_*, is the phase shift between the sources. Unfortunately, due to effect of volume conduction, the phase shift between two signals cannot be directly estimated from the empirical data. Our aim was to reliably estimates the patterns of the spurious amplitude-coupling rather than its absolute value. Therefore, we assumed a constant phase shift of 90 ° for all connections. At 90 °, the absolute value of the spuriously induced amplitude coupling is maximal (Palva et al., 2018). As control analyses, we repeated the simulations for phase shifts of 45 ° and 22.5 °.

We determined the amplitude *A(t)* vectors as follows:

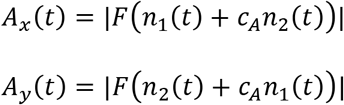

where *n_1_(t)* and *n_2_(t)* are vectors of normally distributed random numbers with data length of 300 s, a pink spectrum and a sampling frequency of 400 Hz approximately matching the original data (Larson-Prior et al., 2013). The || operator refers to the modulus. *c_A_* denotes the ground truth amplitude coupling between the sources *x* and *y* and was set to 0. The function *F* is the complex wavelet transformation of the vectors at the frequency of interest. The wavelet transformation parameters matched our analysis of the empirical data (see above). Analogously, we generated the phase *p(t)* vectors:

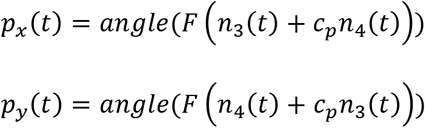

where *n_3_(t)* and *n_4_(t)* are again 300 s vectors of normally distributed random numbers with a pink spectrum at a sampling frequency of 400 Hz. All 4 *n*-vectors were drawn anew for every connection. *c_p_* denotes the ground truth phase-coupling and was set to the empirical weighted phase lag index for every connection, frequency and subject. Finally, we computed the amplitude coupling of the orthogonalized signals *x* and *y* (see above) to quantify the strength of amplitude coupling (*AC_spur_*) that would be expected given the empirical parameter and no ground truth amplitude coupling (*c_a_* = 0). We computed the full correlation matrices for every subject and frequency. We used these matrices to investigate the similarity of spatial patterns derived from the simulations (*AC_spur_*) and the empirically measured amplitude coupling (*AC_meas_*). Importantly, the magnitude of a single *AC_spur_* connection is not informative, because it depends on the empirical phase shift, which is unknown. Thus, under the assumption of a constant phase shift, we correlated the spatial patterns *P_ACspur_* of spurious amplitude coupling with the spatial patterns of empirically measured amplitude coupling *P_ACmeas_*, which discounts magnitude offsets and scaling.

### 4.7 Reliability estimation

To compare the reliability, i.e. reproducibility, of functional connectivity measures, we correlated the seed patterns within (Fig. 3) and between subjects (Fig. 3, 4 & 7). We correlated each column in the correlation matrices pairwise, either between subjects (between-subjects reliability *rel_bs_*) or between recording runs (within-subject reliability *rel_ws_*).

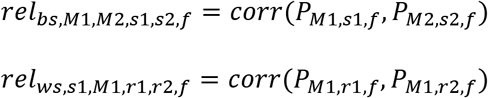

where *M1 and M2* denote the different connectivity measures (*AC_spur_, AC_meas_*, or PC). A pattern *P* describes the connectivity of a given seed with the rest of the source-model, i.e. one column of the full correlation matrix. *M1 and M2* can be the same (within measure) or different (between measure) for the between subject reliability *rel_bs_*. *s*1 and *S*2 denote the subjects involved in the computation, where *s1* ≠ *s2* for the between subject reliability *rel_bs_*. For the within subject reliability, we average the three comparisons (*r_1_* with *r_2_*, *r_1_* with *r_3_*, *r_2_* with *r_3_, r* denotes the run) within a subject. For the between-subject reliability, we first averaged the correlation matrices acquired in the three runs of each subject before correlating between subjects. All reliabilities were independently computed as a function of frequency *f*. We averaged reliabilities across all subject-wise comparisons.

Only reliable signals can be correlated and attenuation correction can only be applied to reliable signals. Therefore, we statistically tested for reliabilities larger than zero (onesided t-test, df = 94 (n = 95), FDR correction, see below) and excluded connections with non-significant reliability.

### 4.8 Pattern similarity, inter-measure correlation and attenuation correction

We correlated the correlation patterns between different metrics, i.e. *AC_spur_* vs. *AC_meas_*, *AC_spur_* vs. *PC, AC_meas_* vs. *PC*:

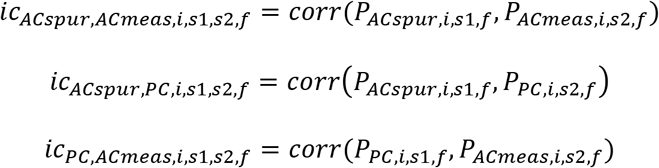

The inter-measure correlation *ic* between two metrics is defined as the Pearson correlation (*corr*) of the seed connectivity-patterns *P* at seed *i* and frequency *f* computed between different subjects *s1 ≠ s2*. The seed pattern *P* is defined as the connectivity of a given seed with the remaining 456 source points, i.e. one column in the full correlation matrix. We computed the inter-measure correlations *ic* for all 8930 unique subject pairings (95^2^-95), 457 connectivity patterns, 3 metric combinations, and 29 frequencies. The measured inter-measure correlations do not only reflect the true underlying similarity of patterns but also the reliability with which these patterns are estimated. Measured correlation decreases with decreasing pattern reliability even if the true underlying pattern correlation remains identical (Supplementary Fig. 1, dashed lines). This effect of reliability is known as attenuated correlations (Spearman, 1904). Following Spearman (1904), we corrected for this attenuation and normalized the mean inter-measure correlation *ic_M1,M2_* by the pooled reliabilities within the measures *rel_M1_* & *rel_M2_*.

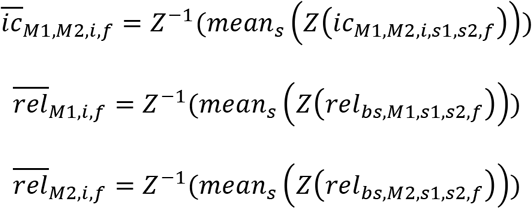

where the mean inter-measure correlation *ic* for different metric combinations *M1/M2* is defined as the mean over all possible subject combinations *s* with *s1 ≠ s2*. The same averaging is done for the reliabilities within the measures *M1* and *M2*. The function *Z* denotes the Fisher Z-transformation and *Z^-1^* the inverse transformation:

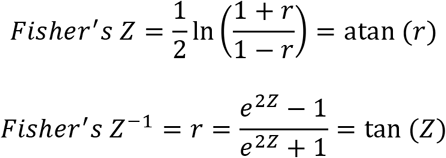

Here, *ln* describes the natural logarithm, *atan* the arcus tangent, *tan* the tangent, *e* Euler’s number and *r* is the correlation coefficient. Finally, the attenuation corrected inter-measure correlation *icc* is defined as:

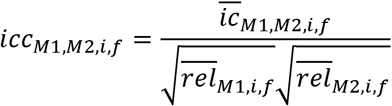

### 4.9 Simulation of attenuation corrected correlations

The attenuation corrected inter-measure correlation is unbiased. I.e., independent of the reliability, with which two patterns are measured, their expected attenuation corrected correlation is the true underlying correlation of these patterns. This is well illustrated by simulations (Fig S1). We simulated two connectivity patterns by drawing (n = 45) from a normal distribution and applying the inverse Fisher’s Z transformation. We controlled the correlation between the two patterns at r = 1 and r = 0.3 as follows: We applied the Cholesky factorization *chol* to the full desired correlation matrix *R* of the two patterns *P_i_* and multiplied the patterns by the resulting matrix:

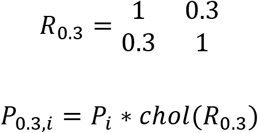

We only used simulated patterns for which |r_P1,P2_ - r| < 0.05. We replicated the patterns for n = 95 subjects and added inverse Fisher’s Z-transformed normally distributed noise independently to every subjects patterns with varying signal-to noise ratios: 0.5, 1, 2. Then, we computed the inter-measure correlation between patterns and the attenuation corrected correlation patterns and repeat the simulation 10,000 times for every SNR. The results are shown in Supplementary Fig. 1.

### 4.10 Statistical testing of attenuation corrected correlations

A perfect attenuation corrected correlation (*icc* = 1) indicates that two cortical patterns are identical if there was perfect reliability. A value smaller than 1 indicates that there is a difference between the two patterns that cannot be explained by reduced reliability. For statistical testing of *icc* against 1, we applied leave-one-out Jackknifing and computed *icc* pseudo-values for each subject, source and frequency. We tested the generated pseudo-value distributions for normality using the Kolmogorov-Smirnov test. We performed one-sided t-tests against 1 when appropriate. We corrected the resulting p-values with false-discovery rate correction within each frequency (Benjamini and Hochberg, 1995).

Notably, statistical testing re-introduces the confound of reliability. While the mean pseudo-values of attenuation corrected correlations are independent of reliability, their variability across subjects increases with decreasing reliability (see also Fig. S1). This confound needs to be taken into account when interpreting the statistical significance.

### 4.11 Quantifying the difference between spurious amplitude-coupling, genuine amplitude-coupling and phase-coupling patterns

We addressed two major questions: First, are there measured amplitude-coupling patterns that cannot be explained by the spurious amplitude-coupling patterns due to phase-coupling? The statistical Null hypothesis for this question is: *icc_ACspur,ACmeas_ = 1*. I.e., the Null-hypothesis is that that the measured and the spurious amplitude-coupling patterns are identical if reliabilities are taken into account. We addressed this question with the methods presented above (Section 4.8 & 4.10). That part of measured amplitude-coupling that cannot be explained by spurious amplitude-coupling we refer to as genuine amplitude coupling (*AC_gen_*). Thus, the quality of the estimated genuine amplitude-coupling depends on the quality of the simulations employed to estimate *AC_spur_*.

The second question was: If there is genuine amplitude coupling, is this amplitude coupling distinct from phase coupling? The statistical Null hypothesis for this question is: *icc_ACgen,PC_ = 1*. I.e., the Null-hypothesis is that that the genuine amplitude-coupling patterns and phase-coupling patterns are identical if reliabilities are taken into account. However, it is impossible to compute *icc_ACgen,PC_* because the genuine amplitude coupling patterns *P_ACgen_* cannot be directly assessed. Yet, it is still possible to test the Null-hypothesis using a geometric heuristic: We conceptualized all connectivity patterns *P* as vectors in a highdimensional space (Fig. 6), where vector co-linearity describes the correlation of patterns. Under the Null hypothesis, the genuine amplitude coupling and phase-coupling patterns are correlated, i.e. *P_ACgen_* and *P_PC_* are co-linear. We assume that the measured amplitude coupling is a summation of the genuine and the spurious amplitude coupling. Thus, under the Null-hypothesis, *P_ACmeas_* lies in the hyper-area between *P_ACspur_* and *P_PC_*. We can now explicate the alternative hypothesis using the co-linearity of these vectors, i.e. their correlation, and the hyper-area between *P_ACspur_* and *P_PC_*: Under condition 1, the correlation between *P_ACspur_* and *P_PC_* is larger than the correlation between *P_ACmeas_* and *P_PC_*. Under condition 2, the correlation between *P_ACspur_* and *P_PC_* is larger than the correlation between *P_ACmeas_* and *P_ACspur_* (Fig. 6C). If one of these two conditions is fulfilled, we reject the Null-hypothesis. Importantly, we used the attenuation corrected correlations *icc_ACspur,PC_,, icc_ACmeas,PC_* and *icc_ACmeas,ACspur_* to test conditions 1 and 2. Thus, the tests are not confounded by pattern reliabilities. We applied leave-one-out Jackknifing to compute three single-subject pseudo-values per source and frequency and performed one-sided paired t-tests to identify sources that fulfill at least one of the two conditions:

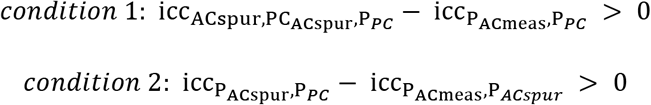

icc_P_ACspur_,P_*PC*__, icc_P_ACmeas_,P_*PC*__, and icc_P_ACmeas_,P_*ACspur*__ refer to the pseudo-value distributions of the correlations at each source and frequency. We conducted a Kolmogorov-Smirnov test of normality on the distributions across subjects and conducted a one-sided Student’s t-test against zero. We use FDR-correction (p < 0.025 corrected) within each frequency to define significance.

### 4.11 Clustering of attenuation corrected patterns

To assess a possible frequency specificity of attenuation corrected similarity between measured and spurious amplitude-coupling patterns, i.e. *P_ACspur_* and *P_ACmeas_*, we clustered the similarity patterns using Gaussian mixed modeling (compare Fig. 5B). This analysis assessed if the differences between the P_ACspur_ and P_ACmeas_ coupling patterns, i.e. icc_ACspur,ACmeas_, are frequency specific. We clustered the frequency specific patterns of similarity using 1 to 8 Gaussians in each model and evaluated the trade-off between explanatory power and complexity of each model with the Bayesian information criterion (BIC). This analysis yielded an optimal model complexity of 4.

We visualized the clustering results in two ways: First, a cross-frequency correlation of icc_ACspur,ACmeas,f_ patterns between all frequency pairs (Fig. 5C). This yielded a frequency-by-frequency correlation between patterns. Second, we used multidimensional scaling (MDS) to represent the patterns in two dimensions (Fig. 5D). We employed MDS based on the pairwise Euclidean distance between all icc_ACspur,ACmeas,f_ patterns.

## Author Contributions

Both authors conceived the study, wrote and revised the paper. M.S.1 performed the data analysis.

## Acknowledgments

The authors declare no conflict of interest. This study was supported by the European Research Council (ERC StG335880) and the Centre for Integrative Neuroscience (DFG, EXC 307). We further thank Constantin von Nicolai, Nima Noury and Andrea Ibarra Chaoul for help with the manuscript and useful discussions. We thank Joerg Hipp for valuable ideas regarding the data analysis.

