## Supplementay Figures for "Dissociated cortical phase- and amplitude-coupling patterns in the human brain"

### Supplementary Figures

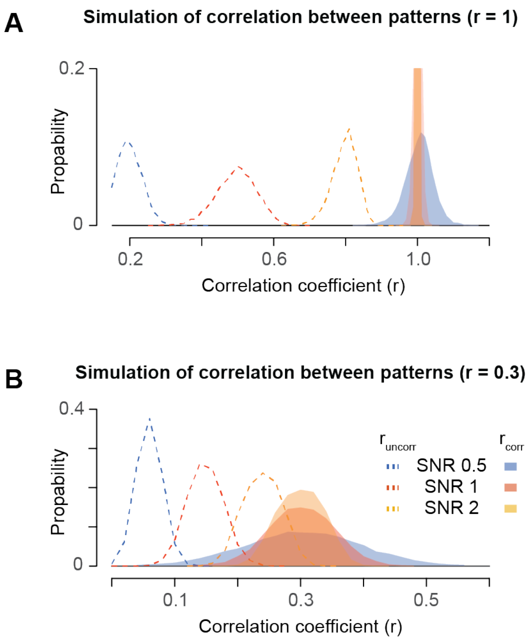


**Figure S1. Simulation of attenuation corrected and uncorrected pattern correlation values for different levels of pattern correlation and between-subject noise**

Distribution of simulated attenuation corrected (solid lines) and uncorrected correlations (dashed lines) between perfectly correlated patterns (A) and patterns correlated at r=0.3 (B). We added uncorrelated noise to the patterns: The signal-to-noise (SNR) levels are set to 0.5, 1 and 2 (for further details see 4.9). The y-axis in (A) is trimmed to 0.2 for visulization purposes: maximum values for SNR 1 = 0.4 and SNR 2 = 0.83.

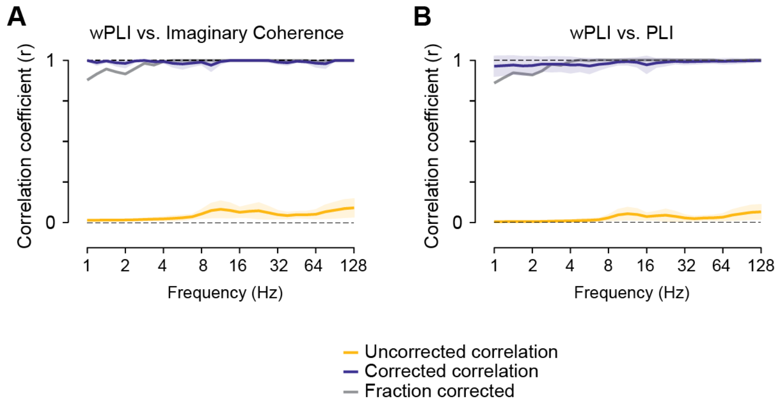


**Figure S2. Comparison between different phase-coupling measures**

Distribution of attenuation corrected (blue) and uncorrected (yellow) correlation between seed-patterns between the weighted phase lag index (wPLI) and imaginary coherency (ImC) (A) and between the wPLI and phase lag index (PLI) (B). The gray line indicates the relative number of corrected sources. Shaded areas indicate the standard deviation across space.

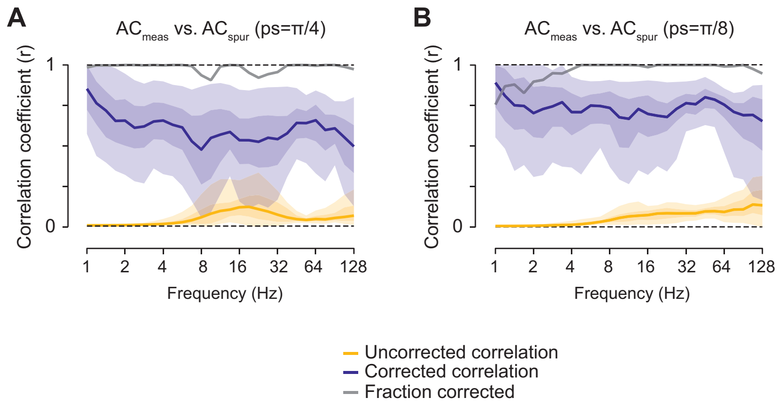


**Figure S3. Correlation between $AC_{spur}$ and $AC_{meas}$ for different levels of phase shift**

Frequency resolved correlation between measured and spurious amplitude-coupling patterns for (A) 45º and (B) 22.5º phase shift between the signals. Lines depict the median corrected (blue) and uncorrected (yellow) correlation. Shaded areas indicate the 5-95% and 25-75% interpercentile range over space.

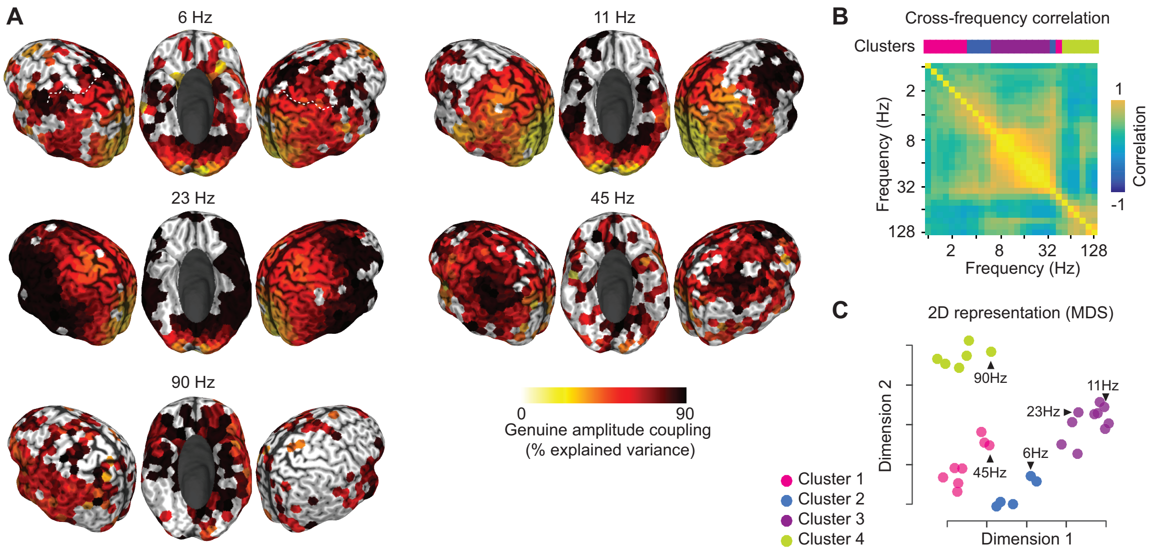


**Figure 5. Spatial distribution of genuine amplitude coupling with statistical masking**

(A) Spectrally and spatially resolved portion of variance in the measured amplitude coupling patterns that cannot be explained by the spurious amplitude-coupling and is thus attributed to genuine amplitude coupling (attenuation corrected). The color scale indicates the amount of genuine variance, i.e. $1-r^2$ of the attenuation corrected correlation between measured and spurious amplitude coupling. The genuine variance is shown for those cortical regions that show an attenuation corrected correlation between measured and spurious amplitude-coupling significantly smaller than 1 ($p<0.05$, FDR corrected) The white dashed line in the top left panel indicates the central sulcus. (B) Cross-frequency correlation of attenuation corrected correlation patterns. The colored bar on top of the matrix indicates the frequency specific clustering of patterns (Gaussian mixture model with 4 clusters for minimal BIC, see 4.11). (C) 2D multidimensional scaling representation of cortical patterns in (A) based on the Euclidean distance between patterns (see 4.11). Points are colored according to the clustering of patterns.
